# A better design for stratified medicine based on genomic prediction

**DOI:** 10.1101/054494

**Authors:** S. Hong Lee, W.M. Shalanee P. Weerasinghe, Naomi R. Wray, Michael E. Goddard, Julius H.J. Van der Werf

## Abstract

Genomic prediction shows promise for personalised medicine in which diagnosis and treatment are tailored to individuals based on their genetic profiles. Genomic prediction is arguably the greatest need for complex diseases and disorders for which both genetic and non-genetic factors contribute to risk. However, we have no adequate insight of the accuracy of such predictions, and how accuracy may vary between individuals or between populations. In this study, we present a theoretical framework to demonstrate that prediction accuracy can be maximised by targeting more informative individuals in a discovery set with closer relationships with the subjects, making prediction more similar to those in populations with small effective size (*N_e_*). Increase of prediction accuracy from closer relationships is achieved under an additive model and does not rely on any interaction effects (gene × gene, gene × environment or gene × family). Using theory, simulations and real data analyses, we show that the predictive accuracy or the area under the receiver operating characteristic curve (AUC) increased exponentially with decreasing *N_e_*. For example, with a set of realistic parameters (the sample size of discovery set N=3000 and heritability h^2^=0.5), AUC value approached to 0.9 *(N_e_*=100) from 0.6 *(N_e_*=10000), and the top percentile of the estimated genetic profile scores had 23 times higher proportion of cases than the general population (with *N_e_*=100), which increased from 2 times higher proportion of cases (with *N_e_*=10000). This suggests that different interventions in the top percentile risk groups maybe justified (i.e. stratified medicine). In conclusion, it is argued that there is considerable room to increase prediction accuracy for polygenic traits by using an efficient design of a smaller *N_e_* (e.g. a design consisting of closer relationships) so that genomic prediction can be more beneficial in clinical applications in the near future.

## INTRODUCTION

The genomics era has demonstrated the polygenic nature of complex genetic traits, and genomic prediction shows much promise for personalised medicine in which diagnosis and treatment are tailored to individuals based on the profiles recorded in their genome. This creates the opportunity for ‘stratified medicine’^1^ in which individuals are classified into higher and lower risk groups and intervention or treatment relevant sub-categories based on profiles that incorporate information from both genomic and environmental risk factors. The utility of this approach, of course, will depend on the reliability of these risk predictions.

A key feature of risk predictors is that their use does not necessarily require an understanding of the aetiology of disease^1^. Usefulness of such prediction is demonstrated by success in genetic selection programs in animals and plants. Risk prediction in human medicine can also have an important impact even in absence of a full understanding of the underlying biology of diseases and disorders. Aggregate effects from causal variants tagged by single nucleotide polymorphisms (SNPs) across the genome can quantify and assess individual risk for a particular disease or disorder, deemed “genomic prediction”.

Genomic prediction has recently been tested and shown to be promising for diseases of which genetic variance is largely explained by a number of major genes^2–4^. However, for polygenic diseases and disorders caused by numerous genes with small effect, which is the case for most complex traits, the accuracy of genomic prediction has been considered too low to be useful in actual clinical applications^5–9^. Most of these studies employed population-based prediction based on unrelated individuals. Several studies have reported a considerable increase in prediction accuracy when the training data set included individuals that were closely related to target sample, from data on humans^10–13^ as well as from other species^14–16^. Some have argued that the use of close relatives may inflate estimated genetic variance due to common environmental effects, or gene-environment or gene-gene interaction^17–19^, and therefore such effects may also bias genomic risk prediction. However, theoretical work from previous studies^20–22^ has shown that genomic predictions are more accurate in populations of smaller effective size, i.e. where individuals tend to be more closely related. In such cases there are effectively fewer chromosome segments to estimate across the genome, which allows a higher prediction accuracy from the same size of data^20–22^. This suggests that subjects that are closely related could be a valuable resource for genomic risk prediction. For predicting human diseases, the area under the receiver operating characteristic curve (AUC) or odds ratio (OR) of case-control status contrasting the higher or lower risk group is a typical measure of prediction accuracy. However, we have no adequate insight in predicting the improvement in AUC or OR when using more related subjects, and how this accuracy may vary between individuals or between populations.

In this study, we revisit the theory on genomic prediction accuracy as presented previously^20–22^, and derive an improved method linking effective population size (*N_e_*) and effective number of chromosome segments (*M_e_*) to prediction accuracy. We also use simulated as well as real data to demonstrate that prediction accuracy can be increased when predicting from more related subjects. We extend this work to a case-control data set, which is a typical design for human diseases, so that the outcomes of this study are applicable to a clinical program for human diseases.

## RESULTS

### Effective number of chromosome segments and prediction accuracy

We validated the theory about estimating *M_e_* (Eq. (10) and (11) in Methods) using the stochastic coalescence gene-dropping method (see simulation I in Methods). The expected *M_e_* (from Eq. ((10) or (11)) was compared to the estimated *M_e_* from the variation in genomic relationships for the subset between discovery and target samples, i.e. using the elements in the off-diagonal block relating to target × discovery sample, and derived from the simulated genotypes using Eq. (12) (Supplementary Figures 1A-3A). Furthermore, the expected prediction accuracy from theory (Eq. (1)) and the observed accuracy from the simulated genotypes and phenotypes were compared (Supplementary Figures 1B-3B).

The estimated *M_e_* from the genomic relationships (using Eq. (12)) agreed with the expected *M_e_* from Eq. (10) or (11) whether using a small or large sample size (supplementary Figure 1A and 2A). From the estimated *M_e_,* the expected prediction accuracy could be obtained from Eq. (1). The expected prediction accuracy was within the confidence interval of the actual observed prediction accuracy over 100 replicates (Supplementary Figure 1B and 2B). With a larger number of chromosomes the estimated *M_e_* from the GRM was close to the expected *M_e_* from Eq. (10)) and (11) that accounts for the correlation between chromosomes, and the expected prediction accuracy from Eq. (1) coincided with the confidence interval of the observed prediction accuracy over 100 replicates (Supplementary Figure 3).

### Theoretical prediction accuracy in relation to *N_e_* as a key design parameter

We theoretically quantified prediction accuracy. Using the theory (Eq. (2), (10) and (11)) that was validated in simulation I, the prediction accuracy for a quantitative trait was quantified in relation to *N_e_,* using *h^2^*=0.5,30 chromosomes each with a genomic length of *L==1* Morgan and *N*=3000 (number of records for the discovery sample) that mimics a typical GWAS. Figure 1 shows that when *N_e_* was smaller, the correlation between the estimated genetic profile scores and phenotypes for the target samples was increased, approaching the square root of the heritability. With *N_e_*=10,000, this correlation was only 0.18, but the accuracy became larger rapidly with smaller *N_e_.* For example, the correlation was 0.65 with *N_e_*=100.

**Figure 1.**
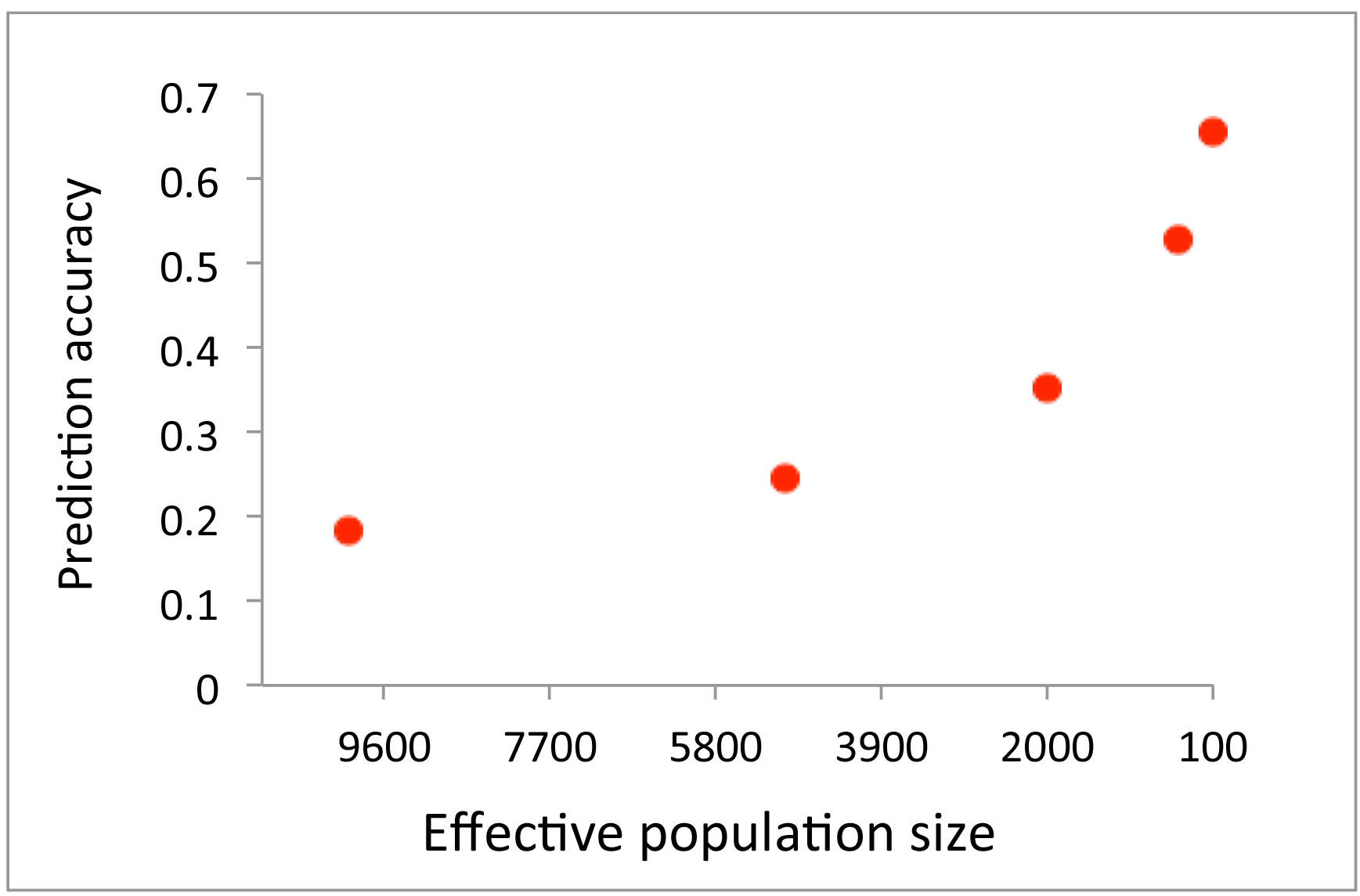
Expected correlation between the phenotypes and estimated genetic profile scores representing the accuracy of genomic prediction of a target sample for quantitative traits when varying *N_e_*=10000, 5000, 2000, 1000 and 100. The number of records (*N*) in the discovery data set is 3000, the true heritability is 0.5 and the number of chromosome is 30 each with a genomic length of 1 Morgan.

The prediction accuracy was also derived for case-control data using the same parameters as above for an underlying quantitative trait. A disease or disorder with population lifetime prevalence of *K*=0.1 and a proportion of cases in the sample of P=0.5 was used. With these parameters, we obtained the expected values for AUC (Eq. (3)), the odds ratio of case-control status contrasting the top and bottom 20% of the genetic profile scores (Eq. (4)) and that contrasting the top 1% of the genetic profile scores and the general population (Eq. (5)). The expected values were verified by comparison with the observed values from simulation II, showing that the expectation and observation were in excellent agreement (Supplementary Figures 4, 5 and 6). Furthermore, we tested the prediction accuracy with a rare disease or disorder with population lifetime prevalence of *K*=0.01, which also showed a good agreement between the expectation and observation (Supplementary Figures 7 and 8).

When using N_e_=10,000, the value for AUC was just 0.60, rising to a value of 0.85 with N_e_=100 (Figure 2). The odds ratio of the case-control status, contrasting the top and bottom 20% according to estimated genetic profile scores, ranged from 2.7 with N_e_=10,000 to 131.9 with N_e_=100 (Figure 3). The odds ratio of the case-control status contrasting the top 1% of estimated genetic profile scores and normal population was 2.3 with N_e_=10,000, and 23.0 with N_e_=100 (Figure 4). With a larger *N* or higher *h^2^*, the prediction accuracy was further dramatically increased (Supplementary Table 4).

**Figure 2.**
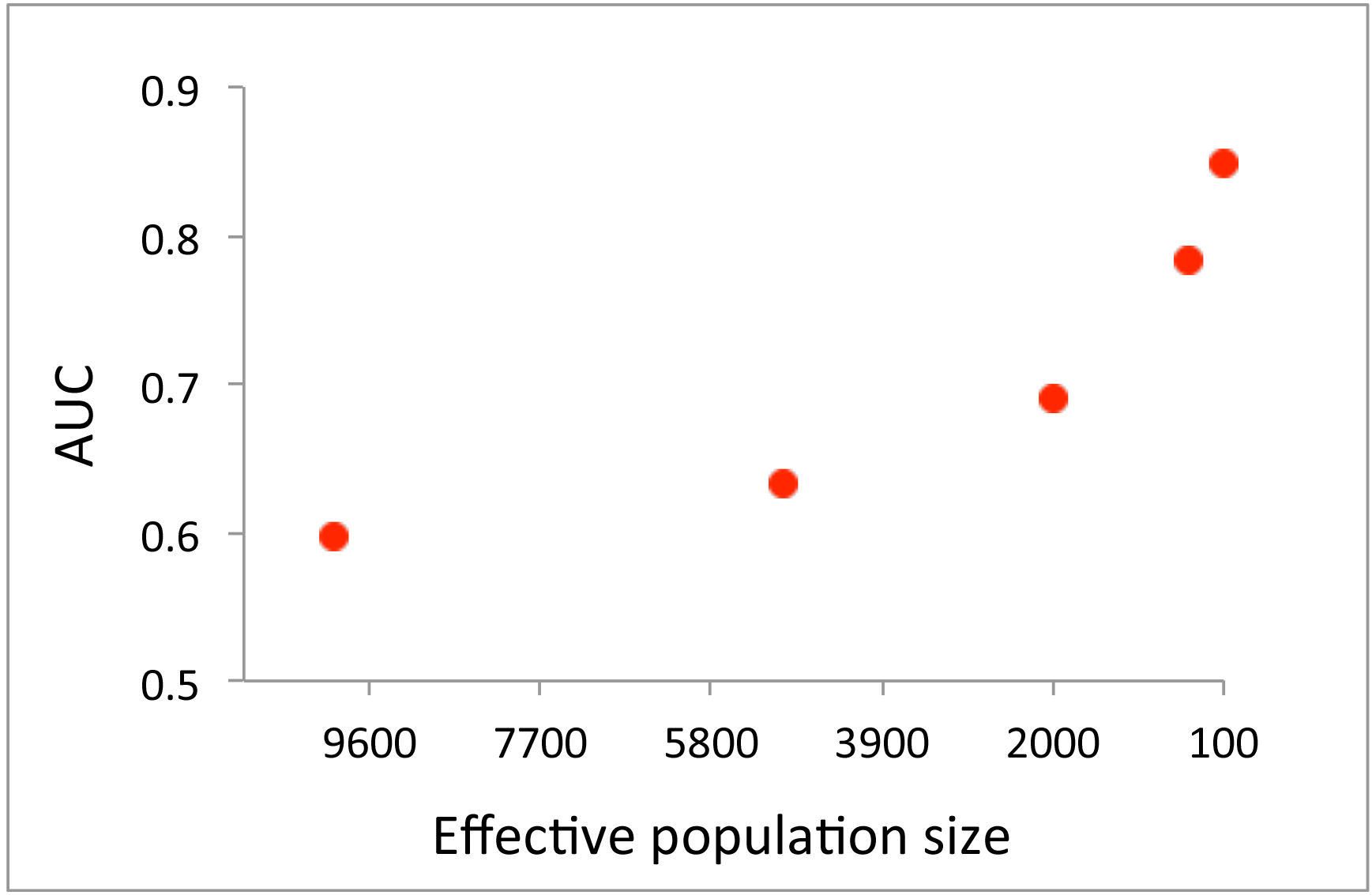
Expected AUC of estimated genetic profile scores in a target sample for case-control data when varying *N_e_*=10000, 5000, 2000, 1000 and 100. The number of records (*N*) is 3000, the true heritability is 0.5, the number of chromosome is 30 each with a genomic length of 1 Morgan, the population prevalence is *K*=0.1 and the proportion of cases in the sample is *P*=0.5.

**Figure 3.**
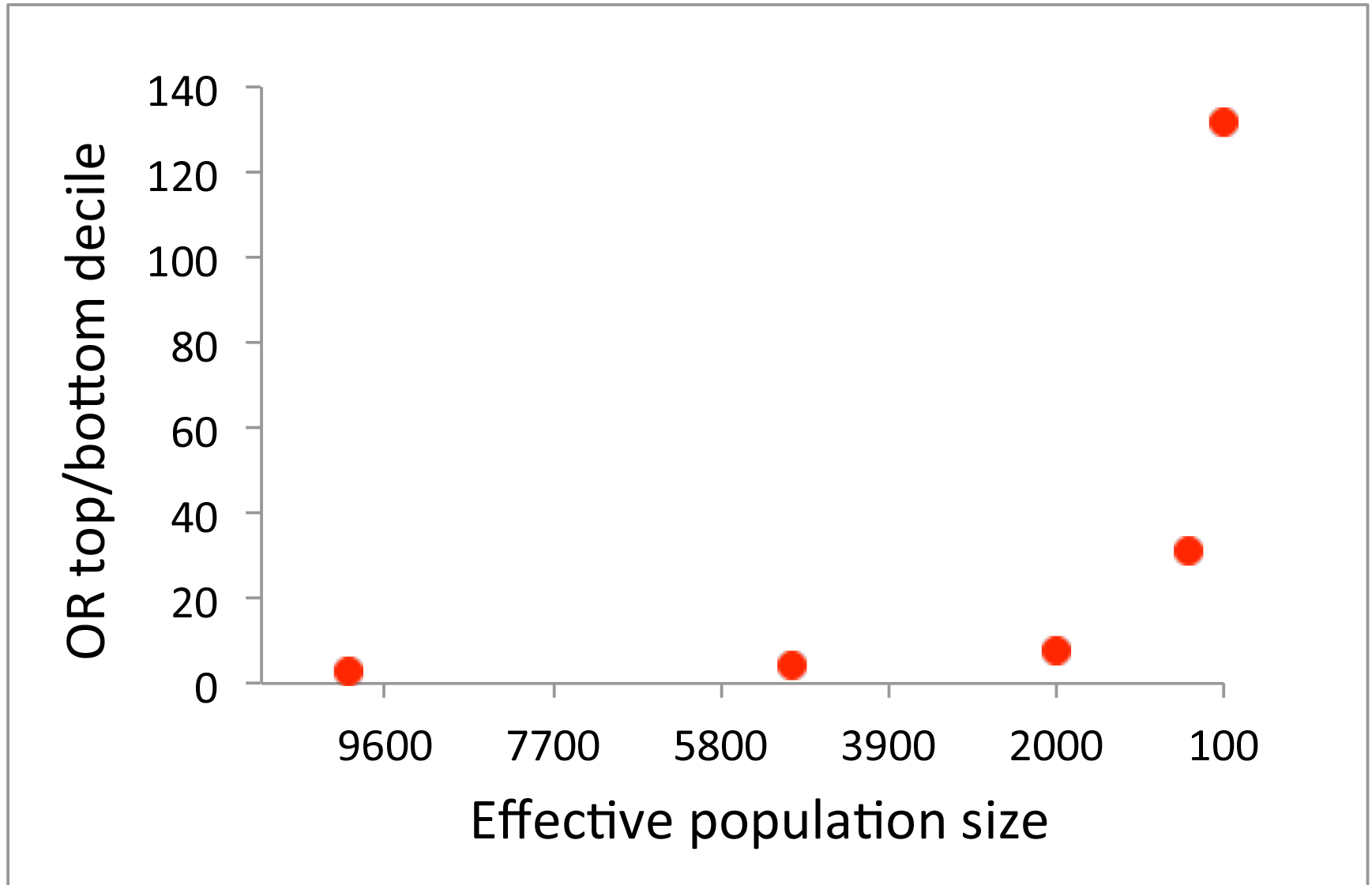
Expected odds ratios of case-control status contrasting the top and bottom 20% of the genetic profile scores in a target sample when varying *N_e_*=10000, 5000, 2000, 1000 and 100. The number of records (*N*) is 3000, the true heritability is 0.5, the number of chromosome is 30 each with a genomic length of 1 Morgan, the population prevalence is *K*=0.1 and the proportion of cases in the sample is *P*=0.5.

**Figure 4.**
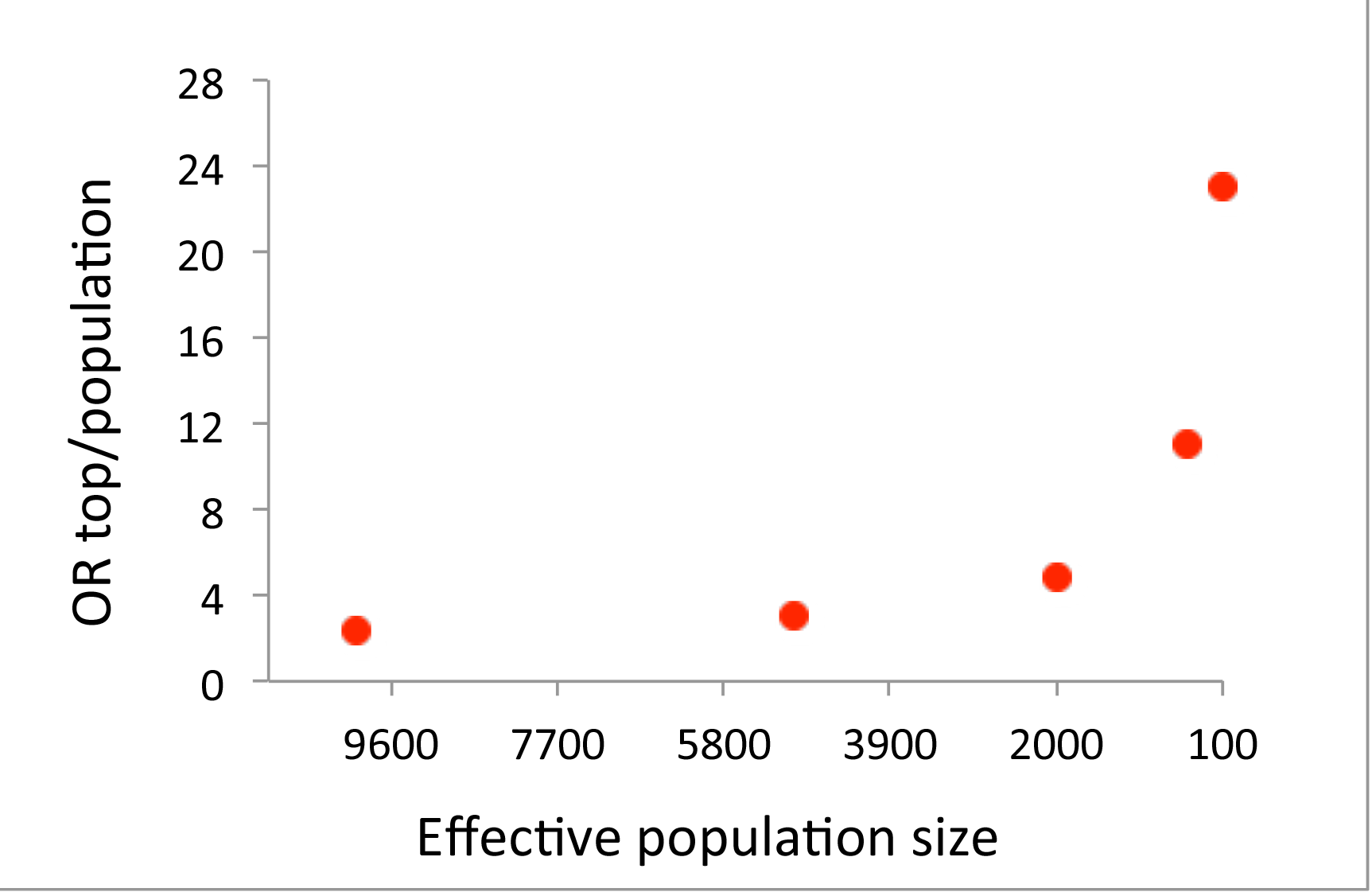
Expected odds ratios of case-control status contrasting the top 1% of the genetic profile scores and a random sample from population in a target sample when varying *N_e_*=10000, 5000, 2000, 1000 and 100. The number of records (*N*) is 3000, the true heritability is 0.5, the number of chromosome is 30 each with a genomic length of 1 Morgan, the population prevalence is *K*=0.1 and the proportion of cases in the sample is *P*=0.5.

### Real data application

We applied the approach to a real data set, the Framingham heart study (see Methods). In 100 cross-validation replicates, the real data was randomly divided into two sets-one for discovery and the other for target, where sampling was either family wise to create a larger *N_e_* or within family to create a low *N_e_.* The discovery set had an average of 3394 individuals and the target set had an average of 849 individuals over 100 cross-validation replicates (Table 1). The estimated *M_e_* from the genomic relationship between the discovery and target samples was 4,434 and 31,080 (from Eq. (12)) when generating a smaller and a larger *N_e_,* respectively. The distribution of variance of relationships, calculated for each target individual when paired with discovery individuals is shown in Supplementary Figure 9 for designs with smaller and larger *N_e_* values. Table 1 shows that the average correlation between the estimated genetic profile scores and the phenotypes (height) in the target set was 0.549 (SD 0.021) and 0.091 (SD 0.043) when using a design with small and large *N_e_,* respectively, clearly indicating the advantage of using a design with a smaller N_e_.

**Table 1.** The accuracy of genomic prediction from a design with smaller or larger N_e_ values when using height phenotypes from the Framingham data.

| | Small $N_e$ | Large $N_e$ |
| --- | --- | --- |
| Quantitative traits (height) - 3394 discovery, 849 target |  |  |
| $M_e$ | 4434 | 31080 |
| Expected accuracy | 0.551<sup>a</sup> | 0.145<sup>b</sup> |
| Observed accuracy | 0.549 (0.021) | 0.091 (0.043) |
| Case-control (10% selection); 680 discovery, 170 target ($K=0.1$ and $P=0.5$) | | |
| $M_e$ | 3247 | 29480 |
| Expected AUC | 0.682<sup>a</sup> | 0.537<sup>b</sup> |
| Observed AUC | 0.687 (0.037) | 0.535 (0.038) |

<sup>a</sup>Expected accuracy from Eq. (2) using the value for $M_e$ and $h^2=0.8^{23-25}$ that is from family studies. <sup>b</sup>Expected accuracy from Eq. (2) using the value for $M_e$ and $h^2=0.45^{26,27}$ that is from population studies. SD over 100 cross-validation replicates is in the bracket.

Interestingly, the results were consistent with the estimated heritability from family-based studies (i.e. h^2^=0.8^23–25^) or population-based studies (i.e. h^2^=0.45^26,27^), which is numerically illustrated in Supplementary Table 5. When using h^2^=0.8 and M_e_=4,434, the expected accuracy of genomic prediction was 0.551 (from Eq. (2)), which was close to the observed accuracy of 0.549 (Table 1). By contrast, using h^2^=0.45 and M_e_=31,080 would give an expected accuracy of genomic prediction of 0. 145, approximately similar to the observed accuracy of 0.091 (Table 1).

Mimicking case-control data, the top 10% of the phenotypes were selected and treated as cases (i.e. K=0.1), and 11.1% of the remaining 90% of phenotypes were chosen to be controls. Therefore, the case-control ratio was 1:1 (i.e. P=0.5). The two sampling strategies used in cross-validation for the quantitative traits, were also used for the case-control data generating higher and lower variance of relationships between discovery and target sets (smaller and larger *N_e_,* respectively/ The discovery set had an average of 680 individuals and the target set had an average of 150 individuals over 100 cross-validation replicates (Table 1). The estimated *M_e_* from the genomic relationship between the discovery and target samples was 3,247 and 29,479 from Eq. (12) for smaller and larger *N_e_,* respectively. In Table 1, the average AUC for the two scenarios was 0.687 (SD 0.037) and 0.535 (SD 0.038), indicating that the AUC was considerably higher with a smaller *N_e_* than with larger *N_e_.* The observed AUC values were very similar to the expected values, based on Eq. (3), for the small *N_e_* design (0.682 withM_e_=3247 and h^2^=0.8^23–25^) and the large *N_e_* design (0.537 with M_e_=29,479 and h^2^=0.45^26,27^), respectively (Supplementary Table 5).

The odds ratio of case-control status comparing each 20 percentile to the bottom 20% of the ranked genetic profile scores demonstrates that the contrasting power was substantially higher with a smaller *N_e_* than with a larger *N_e_.* (Figure 5). The observed odds ratio of case-control status contrasting the top and bottom 20% of the genetic profile scores was similar to the expected odds ratio from Eq. (5) for the small *N_e_* design withM_e_=3,247 and h^2^=0.8^23–25^ and the large *N_e_* design with M_e_=29,479 and h^2^=0.45^26,27^, respectively (Supplementary Table 5).

**Figure 5.**
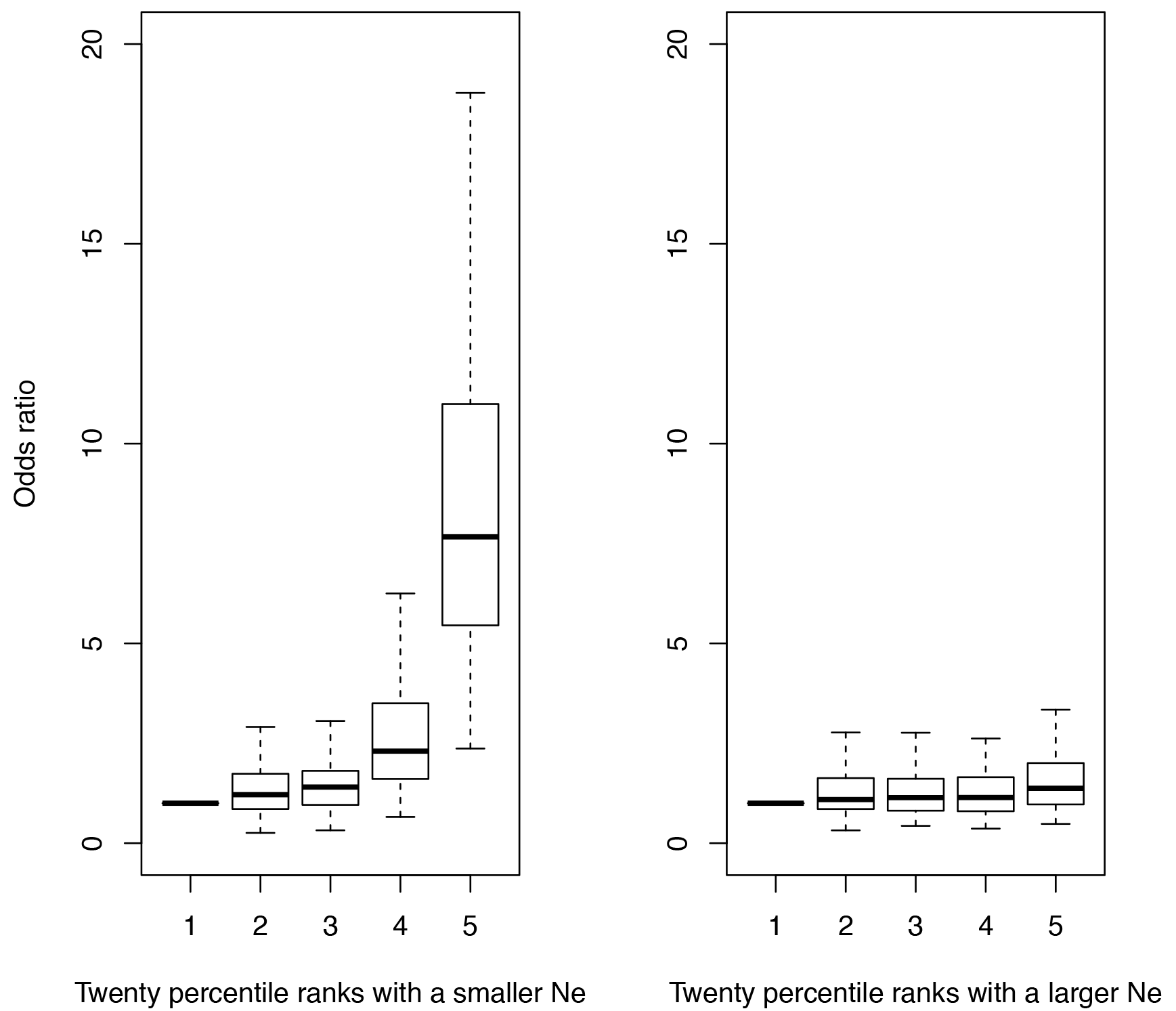
The odds ratio of the case-control status contrasting the top and bottom 20% of the ranked genetic profile scores estimated from a design with a smaller or larger *N_e_*, in the Framingham data

We additionally analysed BMI phenotypes, which gave a similar result in that the prediction accuracy was considerably higher with a smaller *Ne* than with larger *Ne*, and that the observed and expected values agreed with each other (Supplementary Table 6).

When using the GERA dataset that does not have a clear family structure, the prediction accuracy for hypertension phenotypes is significantly higher for 25% of the target sample with the highest variance of pair-wise relationships with the discovery sample (Figure 6 and Supplementary Figure 10). The prediction accuracy was 0.118 (0.114‐0.123) for the top 25% and 0.106 (0.104‐0.107) for the entire target sample. Moreover, the prediction accuracy was significantly decreased to 0.097 (0.095‐0.099) (Figure 7) when higher relationships were removed from the sample (> relatedness of 0.025), therefore increasing *M_e_* (Supplementary Figure 11). These results demonstrate that a higher variance of pair-wise relationships, hence smaller *M_e_*, results in a higher prediction accuracy even when using data from an extensive population-based sample. We also confirmed these results by using the real genotype data (GERA) and simulating phenotypes with the total variance being fully explained by the SNPs in order to support the results from the real data analysis (Figure 6 and Figure 7) by showing that higher prediction accuracy for the top 25% group and the lower accuracy for cut-off high relatedness was not due to non-genetic confounding factors such as artefact batch effects (Supplementary Figure 12 and 13).

**Figure 6.**
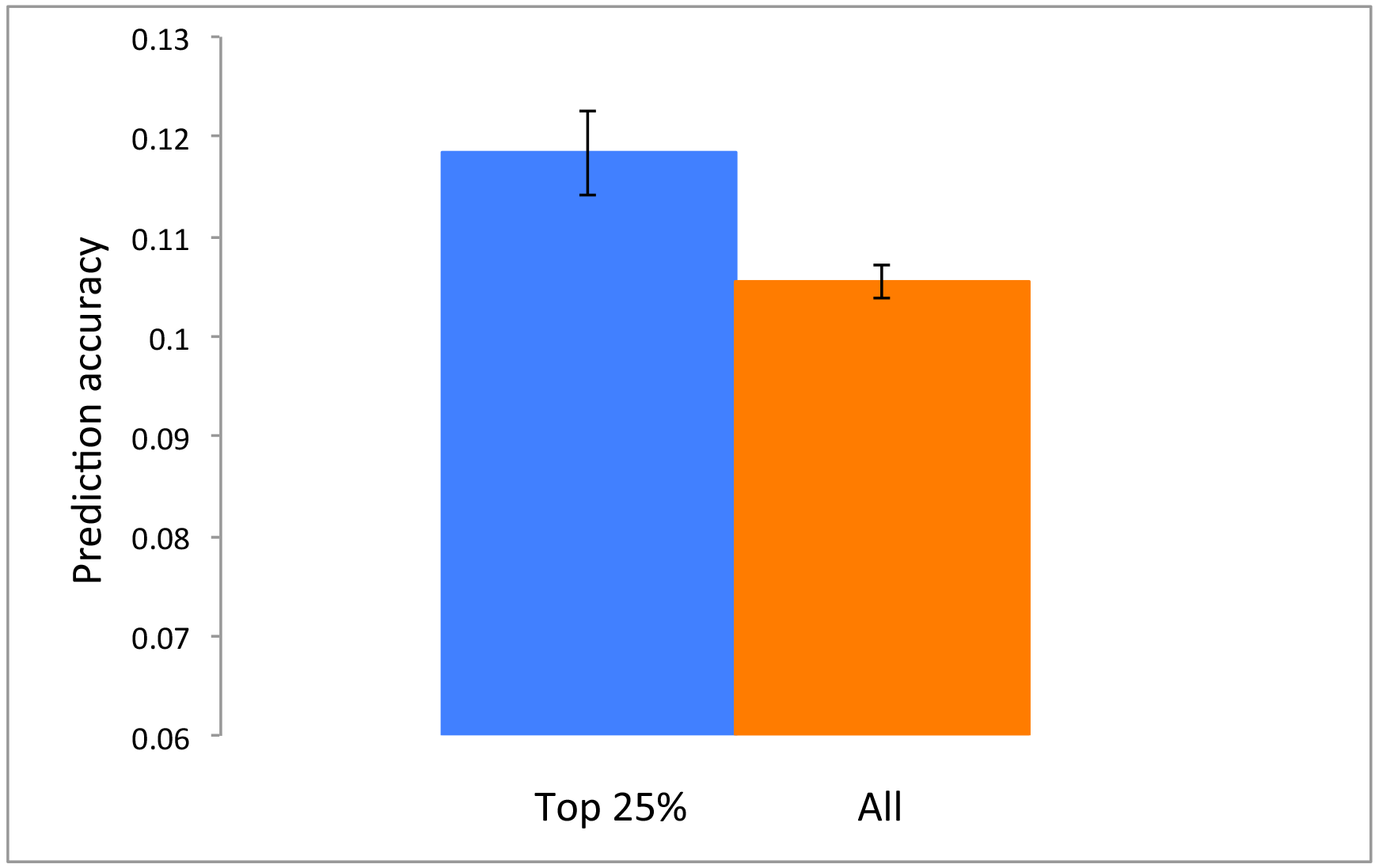
The prediction accuracy is significantly increased when using the top 25% of the target sample according to the variance of pair-wise relationships with the discovery sample (therefore decreasing *M_e_* from 58000 to 37000). GERA data with hypertension phenotypes are used. The error bar is 95% confidence interval of the observed prediction accuracy over 100 replicates.

**Figure 7.**
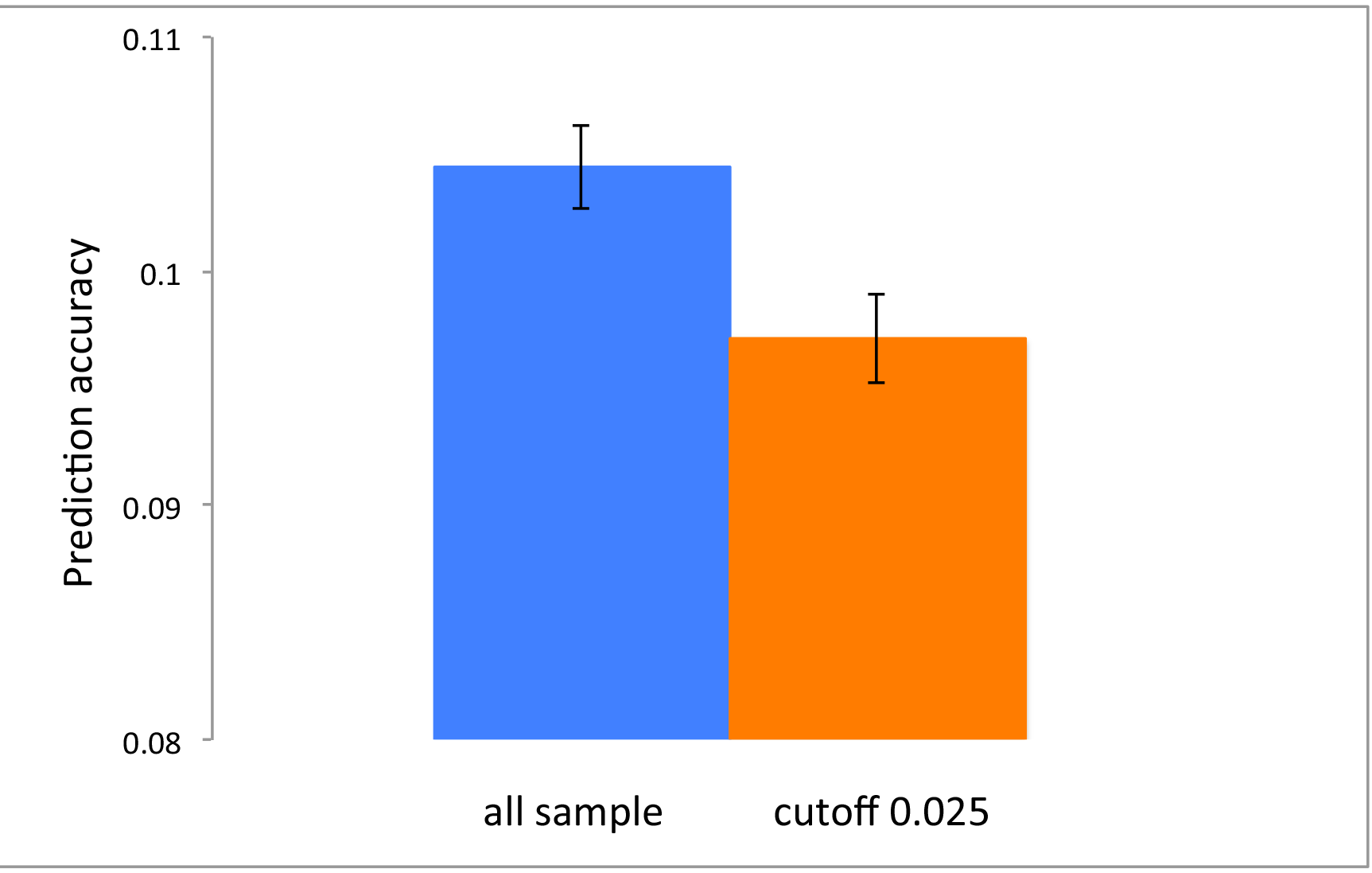
The prediction accuracy is significantly decreased when excluding higher relationships from the sample that results in increasing *M_e_* (from 58000 to 67000). GERA data with hypertension phenotypes are used. The same number of discovery and target sample is used for both tests. The error bar is 95% confidence interval of the observed prediction accuracy over 100 replicates.

We also analysed dyslipidemia phenotypes and found a consistent result showing that the prediction accuracy was significantly increased for 25% of the target sample with the highest variance of pair-wise relationships with the discovery sample (Supplementary Figure 14), and that it was decreased (although non-significant) when higher relationships were removed from the sample (Supplementary Figure 15).

## DISSCUSSION

In this study, we have shown, by simulation and analysis of real data, that genomic prediction that includes closely related individuals leads to higher prediction accuracy. The accuracy can be predicted from the variation in relationships of the target individual with the individuals in the discovery data set. The variation in relationship can be linked back to the number of effective chromosome segments to be estimated, which in turn is a result of a certain effective population size, i.e. the size of a homogeneous unstructured population where the amount of chromosome segment sharing is similar, leading to the same accuracy of prediction. We showed that there is merit in designing the discovery population such that variation of genetic relationships is maximized

Current studies for polygenic diseases or disorders have reported that the accuracy of genomic prediction is not useful for actual clinical practice^5–9^ due to low prediction accuracies. However, it is common practice to use samples from the population that are genetically distant resulting in *N_e_* values of more than a few thousand and a resulting *M_e_* across the genome in the tens of thousands, even when predictions are just within populations of pure European descent. The effective number of chromosome segments is a key parameter on which prediction accuracy depends^20–22^. A desirable design for genomic prediction should have a discovery set that is well related to the target set of individuals, resulting in a smaller *N_e_,* hence lower *M_e_.* It was shown that the prediction accuracy (AUC and ORs) increased with a design of a smaller *N_e_,* compared to that with a larger *N_e_,* using theory, simulations and real data analyses.

The utility of genomic prediction was illustrated with an example where the top percentile of the estimated genetic profile scores had a substantially higher proportion of cases than a random population sample (23-fold) especially when using a design with a smaller *N_e_,* and even when using a moderate sample size in the discovery set (N=3,000) and a heritability of 0.5 (Figure 4). This could be increased to 32-fold with a larger sample size (N=24,000), or 176-fold with a higher heritability (h^2^=0.8) (Supplementary Table 4). This demonstrates that different interventions in the highest risk group could be effectively utilised in a clinical program (i.e. stratified medicine).

Even for a data set of unrelated individuals based on a random population sample, such as the case in the GERA data set, when using the discovery individuals that are more related to the target individuals, the genomic prediction accuracy increased (Figure 6 and 7; Supplementary Figures 12 and 13), because of the larger variance of pair-wise relationships to the target sample (implying lower *M_e_* and N_e_). This may have important implications when only considering population-based samples in genomic risk prediction for human complex traits and diseases.

One challenge with this approach is that a large number of records or samples need to be collected within a local community or from extended families. However, increasingly databases are built with phenotypic and genotypic information from closer relatives^28^. In practice, a composite discovery population combining population-and family-based samples may be an alternative and desirable design, as demonstrated here for the Framingham study as well as in other studies^29,30^. In fact, personalised medicine based on family-based databases are in line with the very concept of family medicine^31,32^.

In many previous studies, it was observed that family-based estimates are considerably higher than population-based estimates for the heritability^25,33,34^. There are plausible explanations for this phenomenon, including inflation due to family effects, gene-gene (G × G) or gene-environment interactions (G × E) ^17–19,35^, or imperfect linkage disequilibrium (LD)^11^ and this has led to many studies discarding information from more closely related individuals. However, the theory and simulation in this study has shown that even in absence of these inflatory effects there will be an increase in prediction accuracy (Figures 1– Figures 4). The results from real data showed that designing the discovery data set to include individuals that are closely related to those in the target sample could give substantially higher prediction accuracy for the target sample. In the real data, this is unlikely to be driven by population stratification, as 10 PCs were included in the analysis model. However, it is possible that non-additive genetic effects could contribute to the increase in prediction accuracy, but one could argue that this is not unwarranted when predicting individual risk. A further study about the possible role of non-additive genetic factors, and whether they can be estimated separately, may be needed.

In the near future, tens of thousands of people will be available for a reference sample to predict genetic risk for a target individual, e.g. all newborn babies could be genotyped and there are improved data bases for recording phenotypes. Using Eq. (1) and (12), we show that either adding more relatives or more genetically distant individuals increased the prediction accuracy substantially (Supplementary Figure 16). The number of relatives required to get the same high accuracy is much lower than that of distant individuals, implying that the information from relatives is of much higher value in an efficient design.

When using case-control data by selecting 10% of the highest phenotypes, the estimated *M_e_* was slightly reduced (Table 1). Specifically, *M_e_* was diminished from 4,434 to 3,247 with a smaller *N_e_* and from 31,080 to 29,479 with a larger *N_e_*. This would be expected because selection on the heritable traits might lower *N_e_^36^*, therefore *M_e_* was therefore decreased.

In this study, it is argued that there is considerable room to increase prediction accuracy for polygenic phenotypes so that genomic prediction can be useful for clinical applications in the near future.

## METHODS

### Accuracy of genomic prediction

Genomic prediction uses phenotypes alongside genome-wide SNPs or sequence data to estimate the effects of observed variants that are projected onto independent subjects and to estimate the subjects' individual genetic profile scores (i.e. breeding values in the context of animal and plant breeding). The accuracy of the genomic prediction depends on the captured genetic variance as a proportion of the total variance, the number of phenotypic observations and the number of independent genomic regions expressed as the effective number of chromosome segments^20–22^, that is

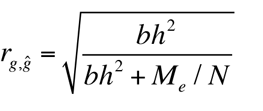

where *r_gĝ_* is the correlation coefficient between the true and estimated genetic profile scores, *h^2^* is the heritability of the trait, *M_e_* is the effective number of chromosome segments, *N*is the number of phenotypic observations and *b* is the proportion of genetic variance captured by observed variants (e.g. SNPs) that can be written as^20–22^

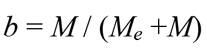

where *M* is the number of observed variants. Owing to dense SNP genotypes or sequence data available, *b* is often close to unity. Therefore, with dense markers, the genomic prediction accuracy can be rewritten as^37^

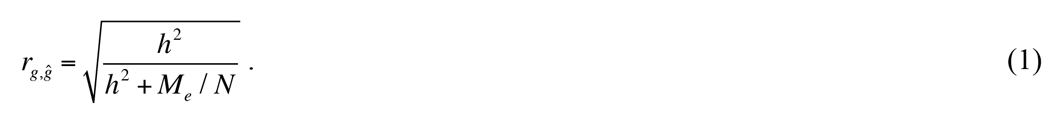

The correlation coefficient between phenotypes and estimated genetic profile scores in a target data set is

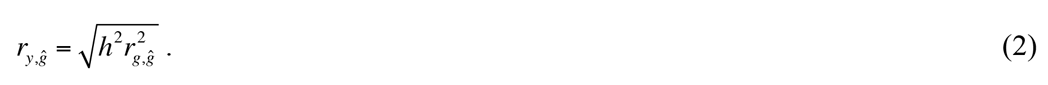

When using case-control studies for human diseases, the correlation coefficient between true and estimated genetic profile scores can be written as^38,39^

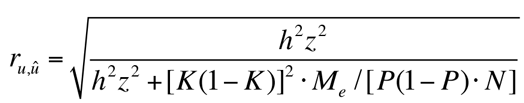

where *u* is a genetic profile score on the 0,1 disease scale^38,40^, *K* is the population lifetime prevalence for the disease, *P* is the proportion of cases in the total sample *N* of cases and controls, and *z* is the height at the threshold on the normal distribution that truncates the proportion of disease prevalence *K* in the liability threshold model. The AUC as a measure of the accuracy of genomic prediction in a target data set for case-control studies is^41,42^

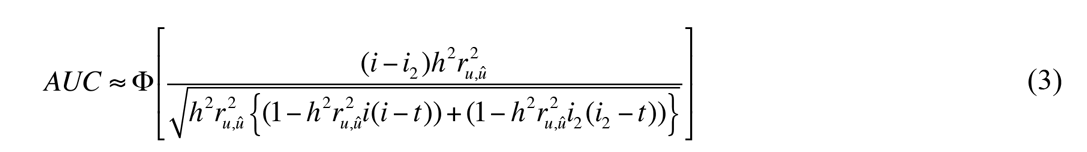

where *i* is the mean liability for cases, *i_2_* is the mean liability for controls, *t* is the threshold on the normal distribution that truncates the proportion of disease prevalence *K* and O is the cumulative density function of the normal distribution.

Another approach to assess the predictive utility of a continuous risk score of diseases, which is common in epidemiology studies, is to stratify individuals into percentiles according to ranked values of the genetic profile scores and estimate the odds ratio of case-control status by contrasting the top percentile with the bottom percentile^5^, that is

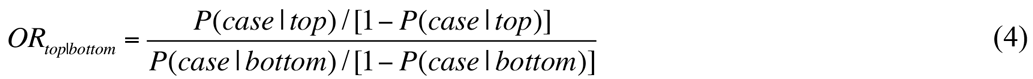

where the probability of being a case in the top/bottom percentile is

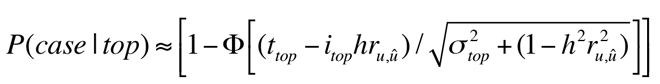

and

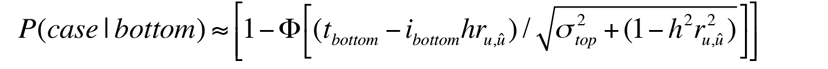

where *i_top_* and *i_bottom_* are the mean genetic profile scores for the top and bottom percentiles, respectively, *t_top_* and *t_bottom_* are the thresholds on the normal distribution
that truncates the proportion of the top and bottom percentiles (for detailed derivation, see Supplementary Note).

In more general terms, it is of interest to obtain the odds ratio of case-control status by
contrasting the top percentile against the general population, that is

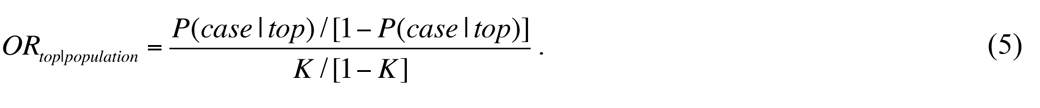

### Effective number of chromosome segments

The effective number of chromosome segments (*M_e_*) is a key parameter for
determining the accuracy of the genomic prediction as fewer segments require fewer
independent parameters to be estimated from the same data, i.e. a higher accuracy. Me
depends on the effective population size (*N_e_*) and the length of genomic region (*L*)^20–22^. There are several studies that derive *M_e_* based on population parameters but there are some inconsistencies between these^20–22^. We revisit the theory and provide another derivation of *M_e_* as a function of *N_e_* and *L*, using the theory of SNP squared correlation matrix^43^.

Considering a genomic region spaning 1 Morgan with M SNPs that are equally distributed over the region, one can construct an M × M squared correlation matrix S in which the elements are the squared correlation coefficients between each pair of SNPs (r^2^)^43^. The squared correlation coefficients can be approximated as *r^2^*=1 / (1+*4N_e_ x c)* where *N_e_* is the effective population size and *c* is the distance in Morgan between each pair of SNPs^44^. Unless the off-diagonal elements in S are all zero, the effective number of SNPs (or chromosome segments) is less than *M.* In order to obtain the effective number of SNPs, each SNP can be weighted and the weights can be obtained as^43^

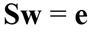

and

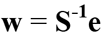

where **w** is an *M* vector of SNP weights derived from the correlation structure among the SNPs and **e** is a vector of length *M* with all elements equal to one. In fact, the effective number of chromosome segments is calculated from the sum of the SNP weights as

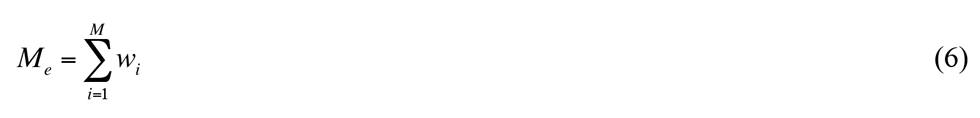

The underlying linear system of order *M* can be written as

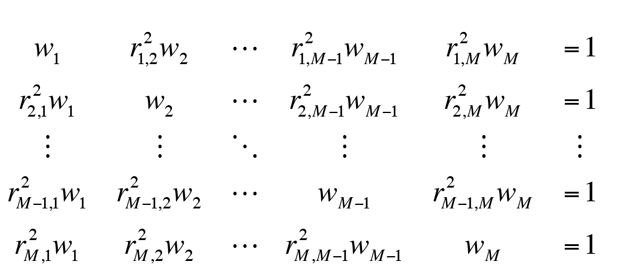

where 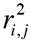 is a correlation coefficient between the *i*th and *j*th SNPs in the matrix **S**.

This linear system can be simplified as

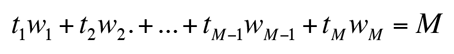

where

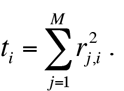

As *t* and *w* are independent random variables, it can be approximated as

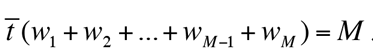

From Eq. (6), the term *w*_1_+*w*_2_+…+*w*_*M*−1_+*w*_*M*_ can be replaced with*M_e_*, resulting in

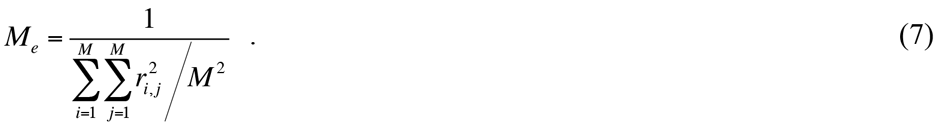

This agrees with Goddard (2009)^20^ who derived this same expression from the covariance statistic between two linked variants while we derived it from the SNP squared correlation matrix theory^43^.

It is noted that the pattern of the same values is repeated in the matrix **S** because of the SNPs being equally distributed. For example, the values for 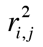 are the same for all combinations for which |i-j| is the same, e.g. 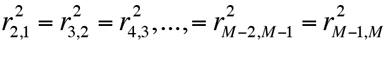. Therefore, the sum in Eq. 7) can be written as

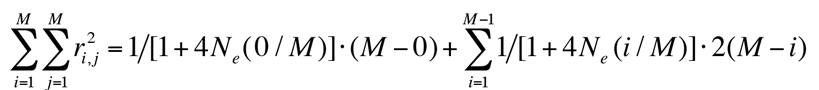

where the first part of each term refers to the estimated *r*^2^ based on the distance, and the second part refers to the frequency of SNP pairs with such an r^2^ value. When scaling the equation by *M*, this can be rewritten as

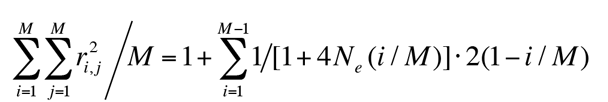

For the right hand side with *M* approaching infinity, the expression can be transformed to a function of *x* having an interval of infinity data points between 0 and 1, which can be written as

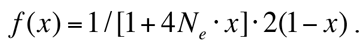

The mean of the function*f(x)* in the variable x ranging from 0 to 1 is defined by an integration. Therefore the denominator in Eq. (7), 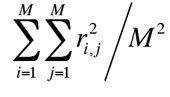 can be obtained as

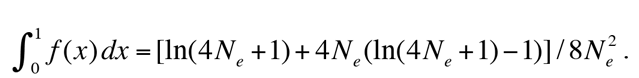

It is straightforward to extend this formula to a genomic region with length *L* rather than 1 Morgan (see Supplementary Note). For an *L* Morgan region, this is

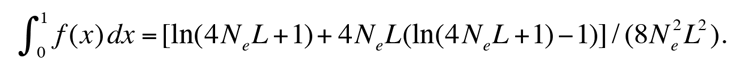

Therefore,

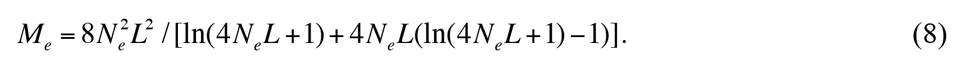

When accounting for mutation^45^, therefore using the correlation coefficients between SNPs from the formula 1 / (2+4*N_e_* x *c*), Eq. (8) is slightly modified to

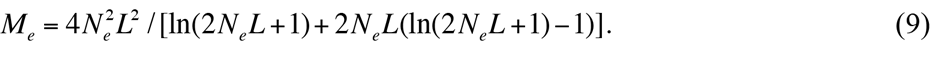

The equivalence between (6) and (7), and the approach linking Eq. (7) and (8) (or (7) and (9)) were validated with actual analyses of the squared correlation matrix (Supplementary Tables 1 and 2).

Eq. (8) and (9) are improved from the previous derivations^20–22^ (Supplementary Table 3). Moreover, previous studies^20–22^ ignored the correlation between chromosomes, however this is not negligible. Following Goddard (2009)^20^ but based on the individual level (rather than the gametic level), the probability of a random pair of individuals having the same parents (i.e. full sibs) in the last generation is (2/N_e_)^2^ and that of having one parent in common (i.e. half sibs) is 4/N_e_-(2/N_e_)^2^. This generates a variance of the relationships among the pairs, which can be analytically approximated as 1/(4N_e_). For the previous generations, the variance is 1/(16*N*_e_), 1/(32*N*_e_), 1/(64*N*_e_) and so on. Summing all these variances gives 1/(3*N*_e_), therefore, the covariance of the pairwise relationships between two chromosomes is 1/(3*N*_e_). Hence, with some rearrangement, Eq. (8) and (9), the expected *M_e_* when accounting for multiple chromosomes, can be expressed as

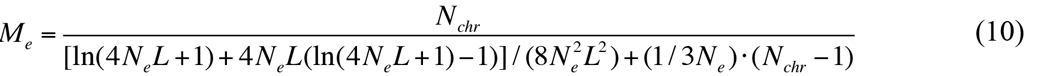

and

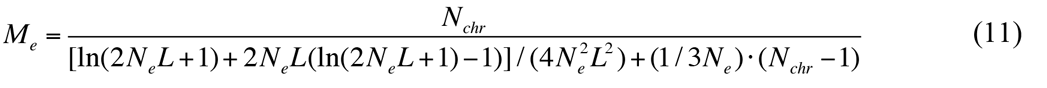

where *N*_chr_ is the number of chromosomes each with length *L*.

### *M*_e_ from the genomic relationship matrix

*M*_e_ can be empirically obtained when a genomic relationship matrix (GRM) is given^21^, which can be interpreted as an observed *M_e_* from the genotype data on which the GRM is based, which can be written as^21^

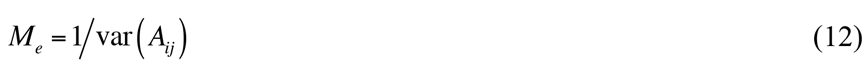

where *A_ij_* is the genomic relationship between individual *i* from the target and *j* from the discovery sample. More details are in the Supplementary Note.

### Estimated genetic profile scores

The MTG2 software^9,46^ was applied to a discovery data set to estimate SNP effects jointly, thereby accounting for linkage disequilibrium (LD) between SNPs. The estimated SNP effects were projected onto the target samples, resulting in a genomic best linear unbiased prediction (GBLUP)^47^ of the genetic profile score for each target individual in the target data set. Dudbridge (2013)^39^showed that the standard genetic profile score method^8,48^ and GBLUP have the same power and accuracy using theory
that assumed all causal variants are unlinked and observed. However, in real situations where there are complex LD structures among SNPs, GBLUP is a preferred method, therefore, we used GBLUP in this study.

### Simulation I

Eq. (10) and (11) were validated with a stochastic coalescence simulation and genomic prediction approach. A stochastic gene-dropping method^49,50^ was used to simulate 20,000 SNPs across a single chromosome of *L*=1 Morgan with *N_e_*=500, 1000, 2000 and 4000 for 500, 1000, 2000 and 4000 generations, respectively. Recombinations occurred across the genomic region according to the genetic distance between SNPs that were equally distributed across the region. The mutation rate was 1e-08 per site per generation^51^. Random mating and selection were used. In the final generation, as a discovery data set, we generated 2000 or 5000 individuals having genotype data for 10,000 causal SNPs, a subset of the 20,000 SNPs, of which the minor allele frequency was more than 1%. For the discovery data set, phenotypes were simulated such that the variance explained by the SNPs was 1% of the total phenotypic variance where SNP and residual effects were drawn from normal distributions. For the target data set, another set of 1000 or 2500 individuals was chosen to estimate the observed accuracy of genomic prediction, i.e. the correlation between true and estimated genetic profile scores. We also conducted simulations of 5 chromosomes each being *L*=1 Morgan long, with a total number of 50, 000 SNPs, resulting in variance explained by the SNPs being 5% of the total phenotypic variance.

Using the genotype data of the discovery data set, a GRM was constructed and *M_e_* was estimated from Eq. (12) as the observed *M_e_* from the simulated data. We used equations (10) and (11) to get the expected *M_e_* given *N_e_* and L. The observed and expected *M_e_* values were compared. In addition, the expected accuracy of the genomic prediction was obtained from equation Equation 1) using the observed *M_e_*, which was compared to the correlation (as the observed accuracy) between true and estimated genetic profile scores (GBLUP) in the target data set.

### Simulation II

In order to confirm the theory in deriving AUC and odds ratios (Eq. (3), (4) and (5)), we simulated disease data (binary responses) using a liability threshold model with a population prevalence of 10% (K=0.1). In the discovery data set, cases were oversampled such that the ratio of cases and controls was equal (P=0.5), mimicking a typical case-control design. The total number of samples in the discovery set was N=3000. We simulated*M_e_* independent SNPs, the effects of which were normally distributed, and a residual effect such that the heritability on the liability scale was h^2^=0.5. We used 5 different values for *M_e_*=254, 1188, 4506, 10891 and 21248, reflecting the expected values for *N_e_*=100, 500, 2000, 5000 and 10000, respectively when using a genomic length of 30 Morgan (30 chromosomes each with 1 Morgan long) and the coalescence formula 1 / (2+*4N_e_* × *c*) (Eq. (9)). SNP effects were estimated in the discovery data set and these estimates were used to predict genetic profile scores in an independent population sample of N=30000 as the target data set. For the target sample, we used a large population sample to reduce empirical sampling error and mimic a realistic situation, e.g. screening newborn babies. We obtained AUC from the genetic profile scores predicting the disease status in the target data set. Additionally, we obtained the odds ratio contrasting the top and bottom 20% of the normal population sample according to the genetic profile scores.

We also obtained the odds ratio contrasting the top 1% according to the genetic profile scores and the normal population. These observed AUC and odds ratios from the simulated data were compared to the expected values from the theory (Eq. (3), (4) and (5)).

## Real data

### Framingham heart study

Publicly available data from the Framingham heart study (phs000007.v26.p10.c1)^52^ were used. Stringent quality control (QC) was applied to the available genotypes, including SNP call rate>0.95, individual call rate>0.95, HWE p-value>0.0001, MAF>0.01 and individual population outliers<6 SD from the first and second principal components (PC). After QC, 6920 individuals and 389,265 SNPs remained. Among them, 4243 individuals from 628 families were phenotyped for height and body mass index (BMI). The mean number of members per family was 6.76 (SD 12.77). Phenotypes were adjusted for birth year, sex, and the first 10 PCs. We calculated the ancestry PCs from the POPRES reference sample^53–55^ because direct PC analysis on the sample could be confounded with family structure^54,56^

In order to contrast designs with smaller and larger *Ne* (and *Me*) values, two kinds of cross-validation schemes were implemented. For a design with larger *N_e_,* 80% of the 628 families were selected as the discovery data set, with the remaining 20% of families used as the target data set. Therefore, the discovery and target sample shared distant common ancestors, hence a larger *N_e_* and *M_e_.* In contrast, for a design with smaller *N_e_,* each member in every family was assigned an 80% chance to be a discovery sample and the rest was in the target sample. Therefore, the discovery and target sample shared recent common ancestors, hence a smaller *N_e_* and *M_e_*.

Using the real genotype data, the genomic relationships between the discovery and target sample were constructed, and *M_e_* was estimated from Eq. (11). The correlation between the phenotypes (that were not used in the analyses) and estimated genetic profile scores in the target data set was estimated.

### Genetic epidemiology research on adult health and aging cohort

As a second real data set, we used genetic data from the European ancestry participants of the Kaiser Permanente genetic epidemiology research on adult health and aging (GERA) cohort^57^, an extensive population sample. The same QC process was applied to the available genotypes. After QC, 62,318 individuals each with 575,760 SNP genotypes remained. We used the trait “hypertension” and “dyslipidemia” for the prediction analyses. Phenotypes were adjusted for birth year, sex, and the first 10 PCs that were inferred from the POPRES reference sample^53–55^.

Unlike the Framingham data, GERA does not have an explicit family structure, i.e. the proportion of pair-wise relationship more than 0.2 was only 0.0002%. Therefore, the family-wise cross-validation schemes used in the Framingham data could not be used. Instead, we randomly selected 46,000 individuals, and randomly assigned 80% and 20% to a discovery data set (N=36,800) and a target data set (N=9,200), respectively, in 100 cross-validation replicates. We calculated the variance of pairwise relationships with the individuals in the discovery data set for each individual in the target data set, and identified the top 25% of the target individuals with the highest variance of the relationships. Then, we tested if the prediction accuracy for the top group (N=2,300) was higher than that for the entire target sample, to show if a larger variance, hence smaller *M_e_*, resulted in a higher prediction accuracy even when using a population-based sample without a substantial family structure. In addition, we obtained the prediction accuracy from a subset of the sample that excluded higher relationships (>0.025). We first applied the relatedness cut-off to all individuals, and then selected 46,000 individuals that were subsequently divided into the discovery (N=36,800) and target data sets (N=9,200). It is noted that for each target individual, the variance of pair-wise relationships with the discovery individuals was reduced due to the relatedness cut-off. In any case, we used the same number of discovery samples (N=36,800) in order to have the same power and for fair comparisons.

## SOFTWARE

Theory, simulation models and GBLUP used in this study have been fully implemented in publicly available software, MTG2. https://sites.google.com/site/honglee0707/mtg2

## ACKNOWLEDGEMENTS

This research is supported by the Australian National Health and Medical Research Council (APP1080157), the Australian Research Council (DP160102126) and the Australian Sheep Industry Cooperative Research Centre. We thank Prof. Peter M. Visscher for helpful discussion in general and his valuable contribution in deriving the theory of odds ratio of contrasting case-control status in percentile analyses. The Framingham Heart Study is conducted and supported by the National Heart, Lung, and Blood Institute (NHLBI) in collaboration with Boston University (Contract No. N01-HC-25195). This manuscript was not prepared in collaboration with investigators of the Framingham Heart Study and does not necessarily reflect the opinions or views of the Framingham Heart Study, Boston University, or NHLBI. Funding for SHARe Affymetrix genotyping was provided by NHLBI Contract N02-HL-64278. SHARe Illumina genotyping was provided under an agreement between Illumina and Boston University. GERA data came from a grant, the Resource for Genetic Epidemiology Research in Adult Health and Aging (RC2 AG033067; Schaefer and Risch, PIs) awarded to the Kaiser Permanente Research Program on Genes, Environment, and Health (RPGEH) and the UCSF Institute for Human Genetics. The RPGEH was supported by grants from the Robert Wood Johnson Foundation, the Wayne and Gladys Valley Foundation, the Ellison Medical Foundation, Kaiser Permanente Northern California, and the Kaiser Permanente National and Northern California Community Benefit Programs. The RPGEH and the Resource for Genetic Epidemiology Research in Adult Health and Aging are described in the following publication, Schaefer C, et al., The Kaiser Permanente Research Program on Genes, Environment and Health: Development of a Research Resource in a Multi-Ethnic Health Plan with Electronic Medical Records, In preparation, 2013.

